# Early neural activity changes associated with visual conscious perception

**DOI:** 10.1101/2021.12.08.471137

**Authors:** Aya Khalaf, Sharif I. Kronemer, Kate Christison-Lagay, Hunki Kwon, Jiajia Li, Kun Wu, Hal Blumenfeld

## Abstract

The neural mechanisms of visual conscious perception have been investigated for decades. However, the spatiotemporal dynamics associated with the earliest neural responses following consciously perceived stimuli are still poorly understood. Using a dataset of intracranial EEG recordings, the current study aims to investigate the neural activity changes associated with the earliest stages of visual conscious perception. Subjects (N=10, 1,693 grey matter electrode contacts) completed a continuous performance task in which individual letters were presented in series and subjects were asked to press a button when they saw a target letter. Broadband gamma power (40-115Hz) dynamics were analyzed in comparison to baseline prior to stimulus and contrasted for target trials with button presses and non-target trials without button presses. Regardless of event type, we observed early gamma power changes within 30-150 ms from stimulus onset in a network including increases in bilateral occipital, fusiform, frontal (including frontal eye fields), and medial temporal cortex, increases in left lateral parietal-temporal cortex, and decreases in the right anterior medial occipital cortex. No significant differences were observed between target and non-target stimuli until >150 ms post-stimulus, when we saw greater gamma power increases in left motor and premotor areas, suggesting a possible role of these later signals in perceptual decision making and/or motor responses with the right hand. The early gamma power findings suggest a broadly distributed cortical visual detection network that is engaged at early times tens of milliseconds after signal transduction from the retina.

## 1. Introduction

With myriad sensory inputs, the nervous system requires a mechanism to identify the most salient signals for additional processing and event emergence in the context of conscious perception. Broadly, this process is known as *detection*, which acts in the earliest stages of sensory processing to rapidly identify target inputs and guides the processing of the most relevant sensory stimuli in higher-order sensory and association cortices for subsequent conscious report (Herman et al., 2019). Because speedy detection is essential for timely decision making and behavior, the stimulus detection network should be active beginning at the earliest times after signal transduction from the sensory organs. For example, signal latency between the retina and primary visual cortex is approximately 50 ms, corresponding to the arrival of signals to the visual detection network (Schmolesky et al., 1998; Herman et al., 2019; Li et al., 2019). Non-sensory cortex also contributes to detection because of long-range cortico-cortical and thalamco-cortical projections allowing for rapid access to sensory signals in association cortices. Several human and non-human primate studies identify the prefrontal cortex as a possible contributor in early signal processing. In non-human primate studies, the frontal eye fields (FEF) were found to be involved in stimulus detection within 100 ms from stimulus onset, even showing visual-evoked response that peaked ahead of V2 and V4 (Schmolesky et al., 1998; Thompson and Schall, 1999, 2000; Bichot and Schall, 2002; Gregoriou et al., 2009; Libedinsky and Livingstone, 2011; Bollimunta et al., 2018). Human studies found similar results with early FEF activity within 100ms from stimulus onset and even as early as 50 ms post-stimulus (Blanke et al., 1999; Muggleton et al., 2003; O’Shea et al., 2004; Kirchner et al., 2009). Beyond the FEF, an fMRI study revealed that the inferior frontal cortex plays a critical role in stimulus detection (Weilnhammer et al., 2021).

Other regions have also been linked with detection, particularly the parietal cortex as a top-down attention feedback structure that communicates with the sensory cortices to modulate the neuronal signals to promote higher-order processing (Critchley, 1962; Corbetta and Shulman, 2002; Saalmann et al., 2007; Gregoriou et al., 2009; Bisley and Goldberg, 2010). This is corroborated by clinical instance of spatial neglect resulting from parietal cortex impairment that can be interrupted as the inability to attend to sensory inputs from a neglected field (Parton et al., 2004). Other regions linked to detection include medial temporal lobe and medial frontal cortex (Wang et al., 2018). Together these findings reveal the detection network possibly engages a broad set of sensory and association regions, both in cortical and subcortical structures, all with early signaling properties. However, many of the nuances of the physiology underlying detection remain unknown.

In the current investigation, we study the spatiotemporal dynamics of cortical networks associated with visual stimulus detection using a continuous performance task (CPT) and intracranial EEG (icEEG) recordings. Despite the limitations of heterogeneous coverage and the confounds of a patient population, icEEG is ideal for examining the human detection network because of its anatomical precision, high temporal resolution, and high signal to noise ratio compared to other human brain imaging modalities, all of which are necessary to capture small, rapid and transient electrophysiology (Parvizi and Kastner, 2018). The analyses we implemented focused on investigating the spatiotemporal dynamics of broadband gamma power (40-115 Hz), which have been shown to reflect the activity of the local neuronal population (Mukamel et al., 2005; Manning et al., 2009; Ray and Maunsell, 2011). Such analyses revealed a broad set of regions involved in stimulus detection including bilateral visual, bilateral prefrontal, bilateral medial temporal cortex, and left lateral parietal-temporal cortex.

## 2. Materials and Methods

### 2.1. Participants

Ten right-handed adult subjects (females=6; mean age=37, range 23–54 years old) undergoing craniotomy with intracranial electrodes for seizure monitoring were recruited from the Yale Comprehensive Epilepsy Program. Subjects participated in the study following written informed consent. All research procedures were approved by the local institutional review board (IRB) at Yale University.

The implanted intracranial electrodes included subdural grids, strips, and depth electrodes. Electrode type, number, and placement were determined by the clinical team overseeing each case. Participants were implanted with on average 196 electrodes yielding a total of 1,957 intracranial electrode contacts across all subjects. Through visual inspection of co-registered structural MRI and whole-brain computed tomography (CT) images, a total of 264 electrodes were excluded across all subjects due to localization in white matter. The remaining 1,693 electrodes were located in grey matter and bilaterally distributed across cortical surface and depth sites (Figure 1B).

**Figure 1.**
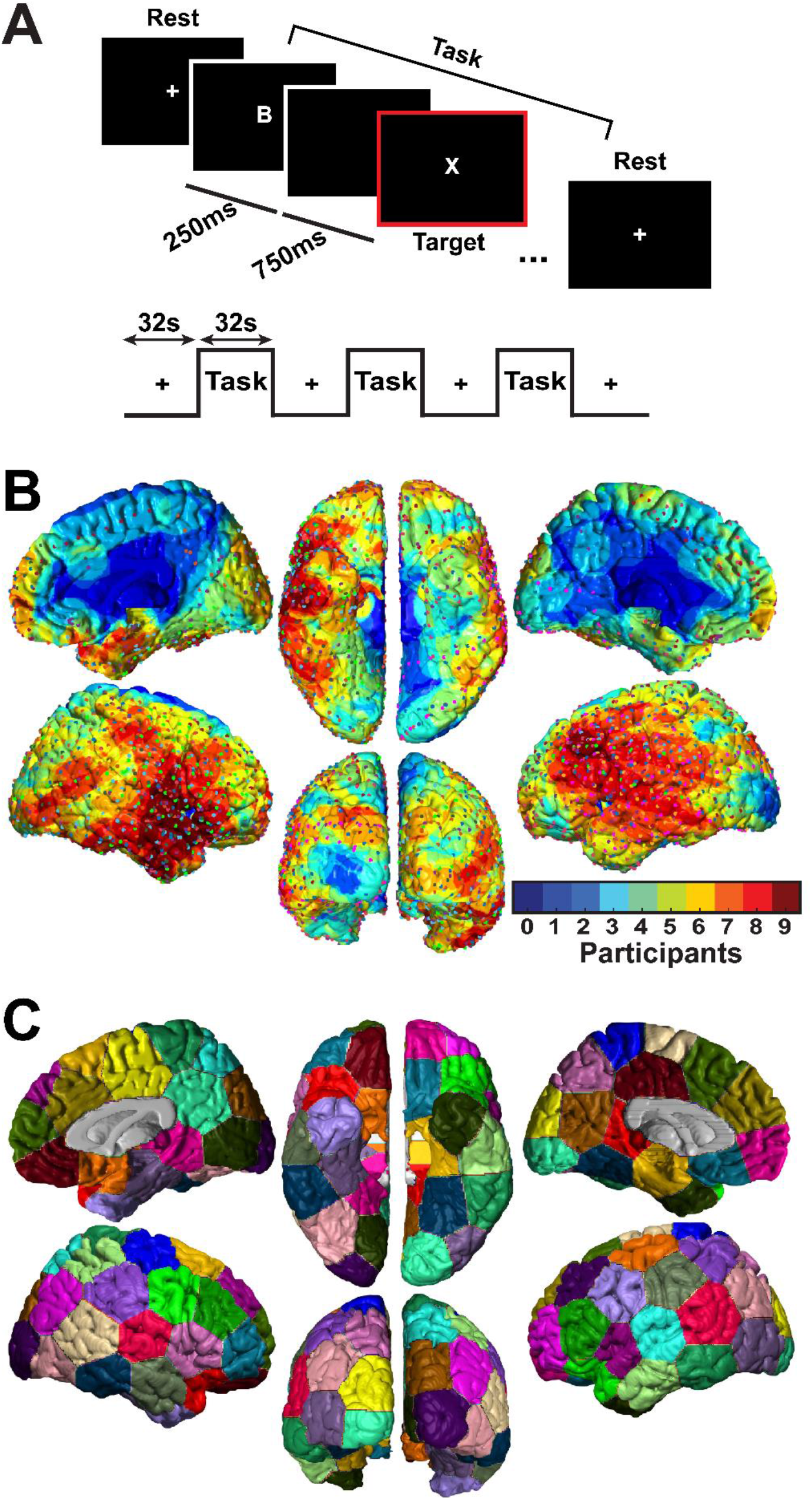
Continuous performance task (CPT), intracranial electrode distribution, and parcellation map. **(A)** CPT task consists of alternating rest and active task phases, each lasting for 32 s. During the active task phase, 32 English letters were presented for 250 ms at a rate of 1 Hz. Participants were instructed to press the right-thumb button whenever a target stimulus (X letter) is presented. **(B)** Density map of 1693 intracranial electrodes implanted in 10 participants is displayed on a common MNI-space brain surface. Signals from each electrode are represented as a spherical space of 15mm radius measured from the electrode central location. The map shows the number of participants contributing to signal analysis at each vertex on the brain surface model. The highest electrode coverage was 9 participants at some brain regions. **(C)** An example of the 120-Parcel bilateral parcellation map was generated using K-mean clustering algorithm. To ensure statistical findings were invariant to changes in parcellation map construction and parcel locations, 100 such parcellation maps were constructed and statistical results were aggregated over the findings from each map (see Methods).

### 2.2. Intracranial EEG data acquisition

The intracranial EEG (icEEG) data were recorded with Natus Neurolink/Braintronics amplifier and pruned using Natus NeuroWorks software (www.neuro.natus.com). EEG data were sampled at 1024Hz and filtered with an analog bandpass filter with corner frequencies of 0.1Hz and 400Hz. The clinical team selected the reference electrode for each participant that best reduced the visible EEG artifacts. The data of one participant were originally sampled at 256Hz. To unify the sampling frequency across subjects, the data of that subject were oversampled using the resample function in MATLAB R2019a (www.mathworks.com) to match the 1024Hz sampling rate of all participants.

To ensure proper synchronization between task event times and the corresponding EEG recordings, transistor-transistor logic (TTL) pulses of varying durations were initiated by the experimental laptop at the onset of task events (button presses, letter presentations, and rest/active phase onsets). The duration of the TTL pulses differed according to the event type to allow differentiation between the events. These TTL pulses were generated by an Arduino Uno (R3; Smart Projects, model A000066) and delivered directly to an open recording channel in the icEEG recording system. The TTL pulses in this channel were used to extract data corresponding to salient task events.

### 2.3. Continuous performance task (CPT)

The continuous performance task (CPT) is a commonly used paradigm to study mechanisms of attention and perception in healthy and clinical populations (Beck et al., 1956; Riccio et al., 2002; Killory et al., 2011). In this study, participants were asked to complete a CPT paradigm coded in Python, run with PsychoPy v1.83 (www.psychopy.org), and presented on a laptop equipped with an NVIDIA GeForce graphics card and 15.6-inch screen display.

The CPT paradigm was composed of alternating blocks of 32-second active task and 32-second rest phases (Figure 1A). During the active task phase, random white capital English letters (A, B, C, D, E, F, G, H, I, L, M, N, O, T, Y, X, and Z; visual angle=1.75°) were presented in random order at the center of the screen on a black background. A total of 32 letters were presented during blocks of the active task phase. The probability of presenting the target letter X within each block was 23.5%. Letters were presented at a rate of 1 Hz and displayed for 250 ms followed by a 750 ms interstimulus interval blank period (black screen) before the presentation of the next letter (or the onset of the rest phase). Participants were instructed to maintain fixation at the center of the screen and immediately respond with a right thumb button press whenever the target letter X appeared. Participants used a response box (In-Line Trainer, Current Designs, Inc., SKU OTR-1×4-L) to deliver their task responses. During the rest phase, participants were instructed to passively view a white fixation cross (visual angle=1.21°) displayed at the center of the screen on a black background.

A single run of the CPT task consisted of 10 blocks of alternating fixation (rest) blocks and active task blocks (5 fixation blocks and 5 active task blocks). The number of runs completed by each participant varied depending on patient in-house availability, duration of stay, and willingness to participate in the study. A total of 63 runs were completed by the 10 participants.

Participants performed the CPT task while being in their hospital bed. The testing laptop was placed 85 cm away from the participant on a bedside table. To ensure having the same ambient lighting conditions across participants and sessions, window blinds were drawn, and lights were switched off during testing.

### 2.4. Epoch segmentation and preprocessing

The event epochs corresponding with letter stimuli presentation were extracted. Each epoch was 4s in duration and centered at the onset of each letter stimulus. The first three letters of each active task block were excluded from the analysis due to a transient signal increase corresponding with the task onset. Letter epochs containing clinical or subclinical epileptic events determined by visual inspection were excluded from the analysis. The remaining epochs were processed using a 4-staged artifact rejection pipeline adopted from Herman et al. (2019) and Li et al. (2019). First, for each epoch, the power spectrum of each electrode was estimated using Welch’s power spectral density method in MATLAB R2019a. To remove trials with high frequency noise, epochs showing peaks with a topographical prominence greater than 200μV/Hz at frequencies greater than 10 Hz were rejected. Next, to detect disconnected or loose electrodes, the mean-square error (MSE) was calculated relative to zero for each electrode within each epoch. Epochs with MSE less than 200 μV^2^ were excluded. Third, the MSE value for each electrode within each epoch was calculated relative to its mean voltage time course averaged within-subject. Epochs deviated from the mean time course by an MSE >3000 μV^2^ were excluded. If more than 20% of the epochs for a subject were marked for exclusion according to this criterion, only the top 20% noisiest epochs would be excluded to balance considerations of noisy data and sample size, as in prior work from our group (Herman et al., 2019; Li et al., 2019). Finally, for each electrode, the standard deviation (SD) was calculated across epochs. Epochs reaching voltages greater than 5 SD were excluded. Across all subjects, this in-house preprocessing pipeline rejected 20% of the letter epochs.

After all letter epochs were passed through the above rejection procedure, the remaining epochs were categorized into specific task event types. Two categories of trials were considered for further analysis in the current study: (1) target letter X trials when the participant successfully responded with a right thumb button press (X-Press trials), and (2) non-target letter trials (all non-X stimuli) when the participant correctly withheld a button response (NonX-NoPress trials).

### 2.5. Broadband z-score gamma power calculation

X-Press and NonX-NoPress trials were filtered using the filtfilt function in MATLAB 2019a to limit the frequency content of these trials within the broadband gamma frequency range (40-115 Hz) as performed in a previous study (Li et al., 2019). The broadband gamma was selected for its known correspondence to population neuronal firing (Mukamel et al., 2005; Manning et al., 2009; Ray and Maunsell, 2011). After filtering, the first and last second of each 4s epoch were removed to avoid filter edge effects. Therefore, the final epoch length was 2s centered at stimulus onset. Gamma power was calculated by squaring the filtered signal and then averaging within 60 ms windows beginning with the first epoch sample, with 30 ms overlap across bins to smooth the power signals over time. The gamma power binning procedure converted each 2s epoch to a total of 65 60ms windows. Trials with smoothed gamma power exceeding 20 SD of the mean power across trials for that electrode within each subject were excluded. The 7 windows representing the time period from -240 to 0 ms corresponded with the pre-stimulus or baseline period while the 32 windows encompassing 0 to 1000 ms post-stimulus were considered as the neural response to the stimulus. The windows representing the time period from -1000 to -240 ms were not considered as a part of the baseline period since they were contaminated by strong neural activity due to preceding stimuli. X-Press trials were z-scored using the mean and SD of the baseline gamma power across all X-Press trials. The same process was applied separately to z-scored NonX-NoPress trials. In particular, for each subject, the gamma power time course at each electrode of each X-Press or NonX-NoPress trial was z-score transformed using the mean and SD of the gamma power during the baseline period of that electrode across all corresponding X-Press or NonX-NoPress trials for that subject.

The z-scored X-Press and NonX-NoPress trials were considered for further examination that employed combined trials (X-Press + NonX-NoPress) and difference of trials (X-Press -NonX-NoPress) analyses. For each subject, combined data were calculated at each electrode by averaging the z-scored gamma power timeseries across all X-Press and NonX-NoPress trials within that electrode regardless of the event type whereas for difference of trials data, z-scored gamma power timeseries were first averaged within each event type for each electrode, and the difference of the electrode average z-scored gamma power was found between the X-Press and NonX-NoPress conditions.

### 2.6. Mapping z-scored gamma power to standard MNI brain

A triangular mesh representing the standard MNI cortical surface was created in BioImage Suite (http://bioimagesuite.yale.edu/) using MNI Colin 22 brain template. BioImage Suite was used to determine electrode locations in MNI space based on each patient’s pre-op MRI, post-op MRI, and post-op CT. Each electrode was assigned to the nearest vertex on the cortical surface. Z-scored gamma power values associated with each electrode were mapped to the standard MNI brain surface as described previously (Herman et al., 2019; Li et al., 2019). In particular, for each participant, all vertices within a 1mm spherical radius from the central vertex to which that electrode was projected were assigned the same z-score values as the central vertex. A linear descending gradient function was used to assign gradually decreasing z-scored gamma values to vertices located within 1-15mm spherical radius from the central vertex where a z-score value of 0 was assigned to vertices at 15mm from the central vertex. Within each subject, the z-score gamma power values associated with each vertex as a result of different electrodes contributing to that vertex were summed. To aggregate the z-scores of each vertex across subjects, a weighted average of these z-scores was obtained across subjects per each vertex by multiplying the average z-scores by the square root of the number of subjects contributing at each vertex location as previously described (Herman et al., 2019; Li et al., 2019).

### 2.7. Statistical analyses

To identify the statistically significant changes in post-stimulus z-scored gamma power compared to the baseline, we employed a spatiotemporal cluster-based permutation test. This approach overcomes the multiple comparisons problem by implementing a single test statistic for the entire spatiotemporal data grid instead of evaluating the statistical significance at each (vertex, time point) pair separately (Maris and Oostenveld, 2007). To reduce dimensionality, the cortical surface mesh was converted to a bilateral parcellation map comprising 120 non-overlapping regions (60 regions per hemisphere) (Figure 1C) generated by applying K-means clustering to the 3-dimensional coordinates of the vertices such that vertices within close proximity were clustered into the same parcel.

The cluster-based permutation statistical approach implemented was adapted from the Mass Univariate ERP Toolbox (Groppe et al., 2011) and applied separately to the combined trials (X-Press + NonX-NoPress) and difference of trials (X-Press -NonX-NoPress). Given that the cluster-based permutation test requires computing an adjacency matrix to identify the neighboring parcels, the test was run for each hemisphere separately. To obtain a cluster-based permutation distribution, the permuted values consisted of mean z-scored gamma power across electrodes within parcels and 60 ms time windows compared to the corresponding baseline values using a paired, two-tailed t-test. For the X-Press + NonX-NoPress analysis, baseline was defined as the (non-zero) mean of the gamma power in the 240 ms prior to the stimulus, whereas for the X-Press – NonX-NPress analysis, baseline was defined as zero, assuming no difference between trials on average prior to the stimulus (this was true in general; data not shown).

For each permutation, prior to calculating t-values, we first randomly shuffled the sign of gamma power values for each electrode to be positive or negative, effectively adding or subtracting it from baseline. This generated a set of t-values across parcels and 60 ms time windows. A parcel and time point were considered eligible to join a cluster based on spatiotemporal adjacency if the t-value fulfilled a set alpha threshold. Positive and negative clusters were defined separately using this threshold. Because the positive and negative values were randomly shuffled, we assumed the permutation distribution to be symmetrical, so to facilitate calculation we only retained negative clusters to create a one-sided distribution. Therefore, the alpha threshold was set at 0.025 (equivalent to 0.05 in a two-sided distribution). Summed t-values for each spatiotemporal cluster were then computed by taking the sum of the t-values for all parcels and time points within each cluster. For each permutation, we retained only the negative cluster with the largest absolute t-value, and collected these across 2000 permutations to create a permutation distribution. To identify significant clusters in the unpermuted data, positive and negative clusters were identified separately, and summed t-values with absolute value above the outer 2.5% of the permutation distribution were considered significant (again equivalent to p<0.05 for a two-sided distribution).

Since the K-means clustering algorithm uses a random seed to initialize the center of the parcels when creating the parcellation map, the algorithm does not converge to the exact same parcels in repeated runs and results in slightly different collections of vertices clustered together within each parcel. In other words, the location of parcels varies slightly across runs of the algorithm. To ensure that the statistical testing was robust and invariant to the changes in parcel locations, we generated 100 separate parcellation maps by running K-means clustering 100 times and we ran the spatiotemporal cluster-based permutation test using each of the generated parcellation maps. This yielded 100 binary significance maps with a value of 1 at the location of significant parcels and a value of zero otherwise. To determine whether a vertex was significant, we used a majority vote approach across the 100 resultant significance maps; vertices that reached significance in more than 50% of the significance maps were considered significant. For both the combined trials (X-Press + NonX-NoPress) and difference of trials (X-Press -NonX-NoPress), we only displayed the z-scored gamma power for vertices that reached significance in more than 50% of the significance maps (see Figures 2, 3).

**Figure 2.**
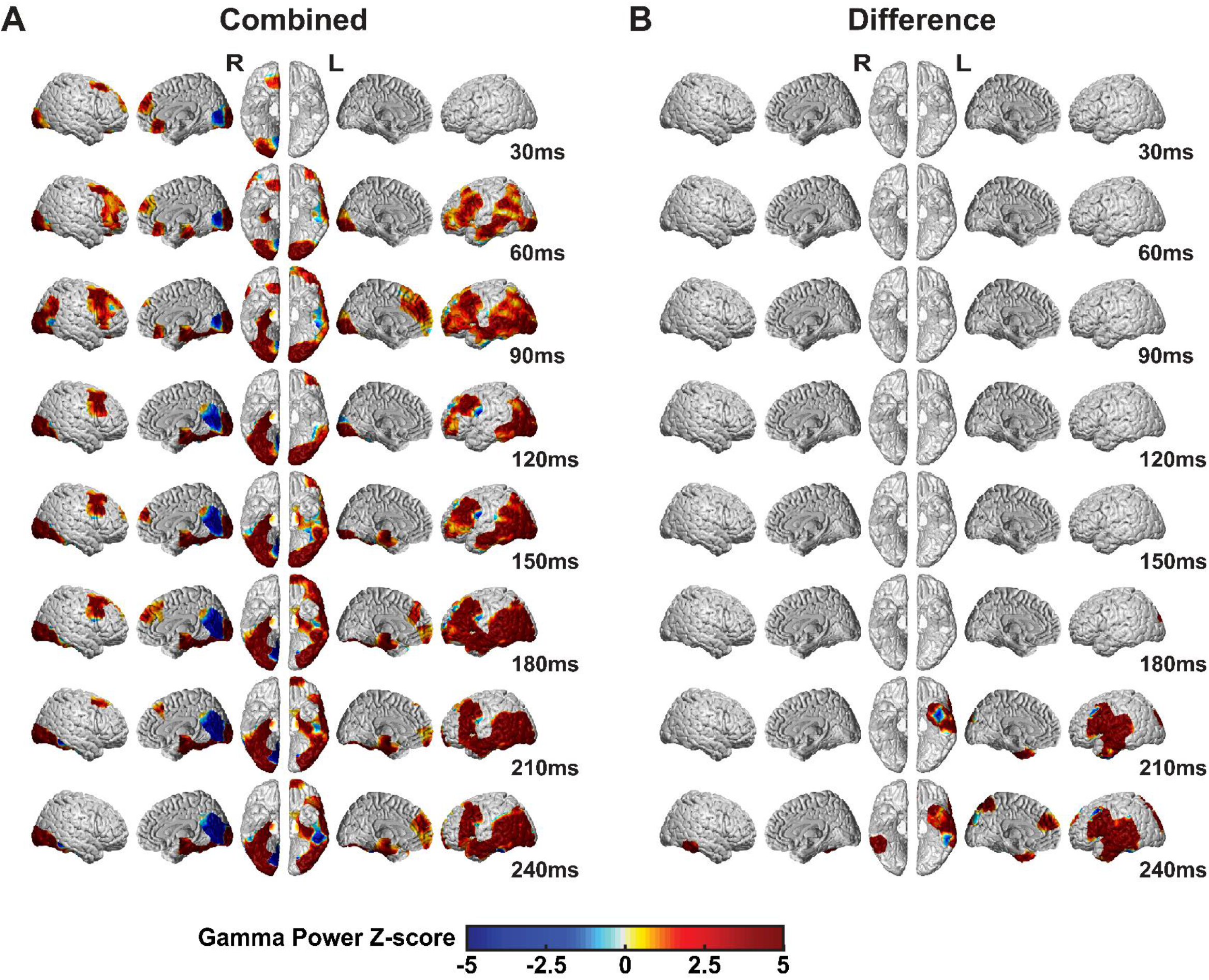
Statistically significant gamma power changes at the early stages of visual conscious perception for combined (X-Press + NonX-NoPress) and difference (X-Press -NonX-NoPress) analyses of trial types (N=10 subjects). The significance was assessed using cluster-based permutation testing (see Methods) and only statistically significant changes are displayed. Displayed times are center points of 60 ms analysis windows. **(A)** X-Press + NonX-NoPress: gamma power increases were observed at the earliest times after stimulus presentation (<150ms) in bilateral visual cortex and fusiform gyrus, bilateral medial temporal parahippocampal gyrus, bilateral caudal middle frontal gyrus overlapping with FEF, bilateral superior frontal gyrus, bilateral orbital frontal cortex, and bilateral inferior frontal cortex. Early increases were also observed in left lateral parietal cortex and left lateral temporal cortex. Decreases were observed in right anterior medial visual cortex **(B)** X-Press -NonX-NoPress: There were no significant gamma power differences at the earliest stage of stimulus presentation (<150ms). At later times differences emerged especially in left cortical areas (right hand was used for button press), seen more fully in Figure 3.

**Figure 3.**
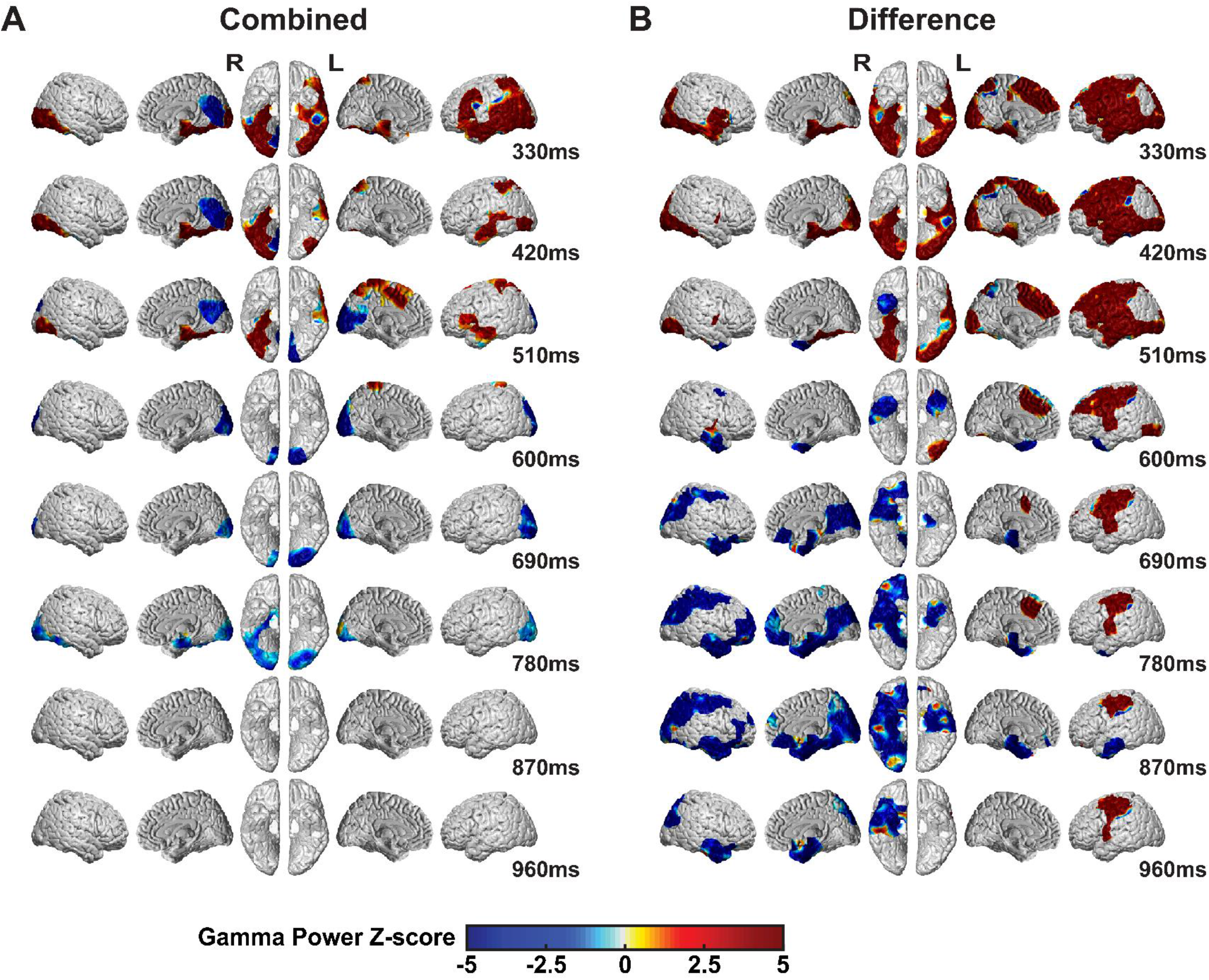
Statistically significant gamma power changes at the late stages of visual conscious perception for combined (X-Press + NonX-NoPress) and difference (X-Press -NonX-NoPress) analyses of trial types (N=10 subjects). The significance was assessed using cluster-based permutation testing (see Methods) and only statistically significant changes are displayed. Displayed times are center points of 60 ms analysis windows. **(A)** X-Press + NonX-NoPress: gamma power increases in the combined data were observed in various brain areas at later times (>300 ms). This wave of increases receded at approximately 510 ms. **(B)** X-Press - NonX-NoPress: gamma power associated with X-Press trials had greater magnitude in multiple regions of the left cortex. At later times, relatively decreased signal for X-Press trials was observed in the bilateral anterior/medial temporal cortex and in wide regions of the right hemisphere.

### 2.8. Regions of Interest

Eight anatomical regions of interest (ROIs) involved in the early stages of visual perception as well as task-specific neural changes were selected. The ROIs were obtained from the SPM12 toolbox (https://www.fil.ion.ucl.ac.uk/spm/) MarsBaR (voxel size=2×2×2mm) in MNI space (Tzourio-Mazoyer et al., 2002) and mapped onto the vertices of the brain surface mesh also in MNI space. The mapping was performed such that any vertex located within a maximum distance of 1.5 mm from the center of an ROI voxel was attributed to that ROI. However, some vertices on the mesh surface were not included in their corresponding ROIs and appeared as holes on the brain surface mesh because they did not fall within 1.5mm from the center of MRI voxels especially at the depths of sulci and at the edges of ROIs so they needed to be corrected manually. Guided by visual inspection, the correction process was performed by changing the maximum allowed distance between the center of a voxel within an ROI and the nearest vertex on the brain mesh until all vertices were assigned appropriately to the correct nearest ROI.

The ROIs (Figure 4A) included occipital cortex (O), fusiform gyrus (FG), parahippocampal gyrus (PH), caudal middle frontal gyrus (CMF), superior frontal gyrus (SF), parietal cortex (P), supplementary motor area (SMA), and precental gyrus (PG). Calcarine, Cuneus, Lingual, Occipital_Sup, Occipital_Mid, Occipital_Inf from MarsBaR were combined to form the occipital cortex (O) ROI. The parietal cortex (P) ROI was formed by grouping MarsBaR’s Parietal_Sup, Angular, Parietal_Inf, and SupraMarginal ROIs. The caudal middle frontal (CMF) ROI is the caudal part of Frontal_Mid from MarsBaR. Finally, fusiform gyrus (FG), parahippocampal gyrus (PH), superior frontal gyrus (SF), supplementary motor area (SMA), and precental Gyrus (PG) ROIs correspond to Fusiform, Parahippocampal, Frontal_Sup_Medial, Supp_Motor_Area, and Precentral ROIs from MarsBaR, respectively.

**Figure 4.**
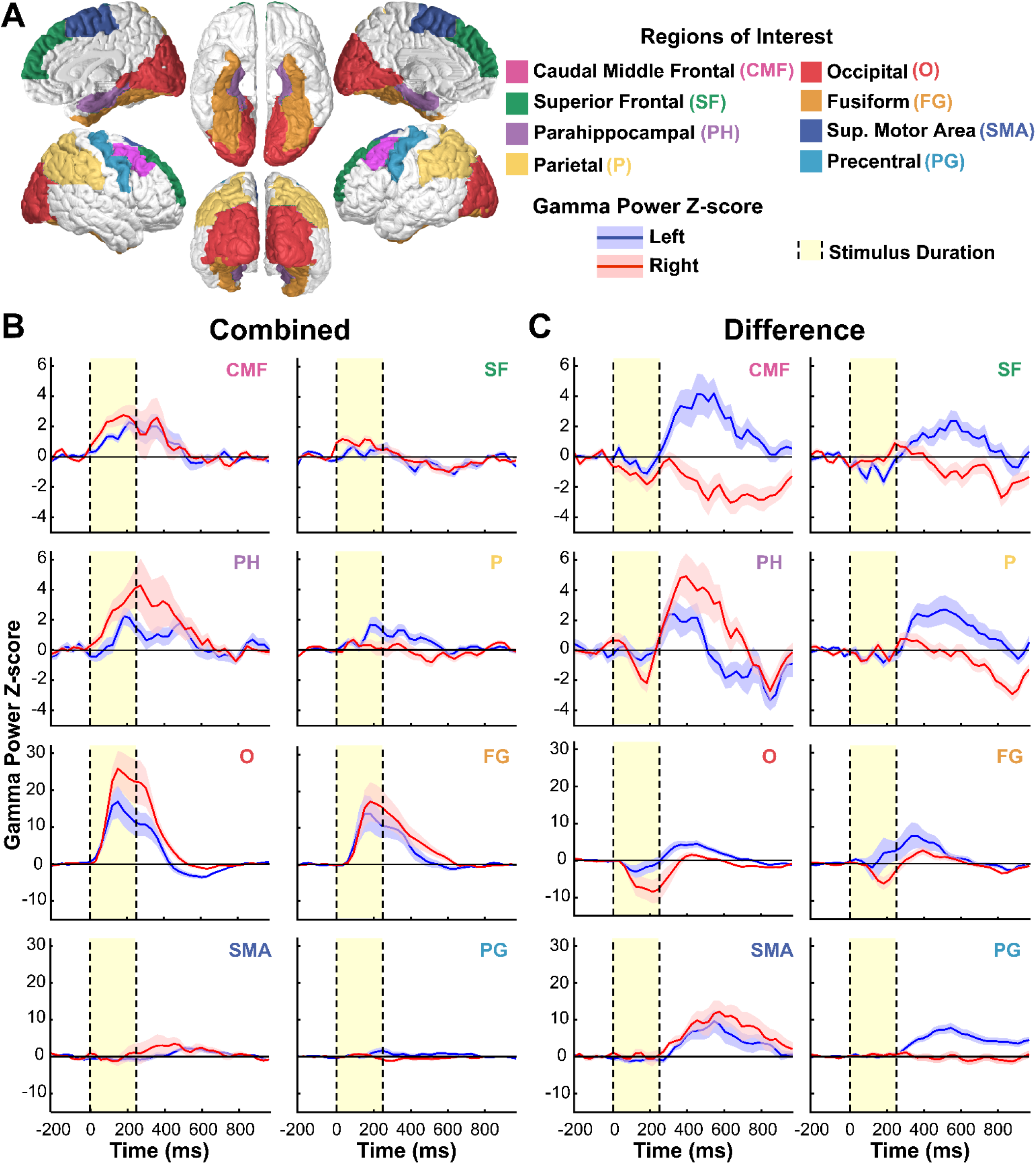
Gamma power time courses of 8 anatomical ROIs. **(A)** Eight representative anatomical regions of interest (ROIs) were identified from the SPM12 toolbox MarsBaR (voxel size=2×2×2mm). The ROIs included caudal middle frontal gyrus (CMF), superior frontal gyrus (SF), parahippocampal gyrus (PH), parietal cortex (P), occipital cortex (O), fusiform gyrus (FG), supplementary motor area (SMA), and precental gyrus (PG). **(B)** X-Press + NonX-NoPress combined analyses: All ROIs except for SMA and PG showed early gamma power increases following the stimulus onset. **(C)** X-Press - NonX-NoPress difference analyses: The most prominent gamma power differences were seen mainly at later times after stimulus presentation. Vertical black dashed lines and yellow shaded regions indicate timing of the 250ms visual stimulus. Note difference in vertical scale in both B and C for the top four traces versus the bottom four traces. Same subjects and data as in Figures 2 and 3.

For each ROI, average combined and difference time courses of the electrodes within that ROI were plotted separately for each hemisphere (Figure 4B, 4C). For every subject, combined data were calculated at each electrode by averaging the z-scored gamma power timeseries across all X-Press and NonX-NoPress trials within that electrode regardless of the event type whereas for difference data, z-scored gamma power timeseries were first averaged within each event type for each electrode, and the difference of the electrode average z-scored gamma power was found between the X-Press and NonX-NoPress conditions. For both combined and difference cases, average across electrodes from all subjects was calculated and reported (Figure 4B, 4C).

## 3. Results

Two categories of trials, X-Press and NonX-NoPress, were analyzed for changes in z-score gamma power following visual letter stimuli presentation. Parallel analyses were applied to (1) combined trials (X-Press + NonX-NoPress) and (2) difference of trials (X-Press - NonX-NoPress) to investigate the similarities and differences between the neural signals resulting from X-Press and NonX-NoPress events, especially at the early times post-stimulus that are hypothesized to be linked to the initial stages of visual signal detection leading to conscious perception. Cortical surface maps displaying gamma power changes are shown starting from the stimulus onset to 270 ms post-stimulus in 60 ms overlapping increments for combined and differences of trials (Figure 2). In Figure 3, cortical surface maps of gamma power changes occurring at later time points from 300-1000ms post-stimulus are presented. Regions of interest (ROIs) and the corresponding gamma power time courses for combined and differences contrasts are shown in Figure 4. The activation onset/offset times reported below are defined as the center points of the 60-ms time windows at which that activation started/ended.

### 3.1. Early gamma power changes

Cluster-based permutation statistical analysis on the difference of trials revealed there were no statistically significant changes between the gamma power responses within the first 150ms post-stimulus between X-Press and NonX-NoPress trial types (Figure 2B). To increase statistical power, we combined the X-Press and NonX-NoPress trials to investigate the shared early changes across the two trial types (Figure 2A).

For combined trials, cluster-based permutation statistical analysis on z-score gamma power changes revealed statistically significant bilateral increases in gamma power in the bilateral primary visual cortex, with an approximate onset of 60 ms post-stimulus (Figure 2A). These changes co-occured with increased bilateral gamma power in the caudal middle frontal gyrus (overlapping with the frontal eye fields) and medial temporal parahippocampal gyrus, with the right parahippocampal gyrus slightly preceding the left. Sustained statistically significant early increases in gamma power were also found in the bilateral superior frontal gyrus (Figure 2A). At ∼60 ms post-stimulus, bilateral orbital frontal cortex and bilateral inferior frontal cortex showed significant gamma power increases (Figure 2A). The right fusiform gyrus showed a prominent gamma power increase at ∼90 ms post-stimulus, while a more gradual spread of increased gamma power across time was observed in the left fusiform gyrus. However, notably, our results regarding the full spatial extent of gamma power increases in the fusiform gyrus are limited by electrode coverage (Figure 1B).

Even at these early time points, not all changes were observed bilaterally: an early sustained gamma power increase (60-330 ms) was observed across left superior and left inferior parietal lobules as well as left lateral temporal cortex. In contrast, early decreases were seen in right anterior medial occipital cortex overlapping with lingual gyrus, cuneus, and calcarine.

The ROI time courses for representative regions correspond with the early signal profiles from whole brain surface gamma power responses reported in Figure 2. Combined time courses showed bilateral increased gamma power in the occipital cortex, fusiform, caudal middle frontal, and parahippocampal gyri within the first 150 ms post-stimulus onset (Figure 4B). Early increases with smaller magnitudes were present bilaterally in superior frontal and parietal ROIs. Notably, the early changes in the above-mentioned regions, except for the parietal ROI, were greater in magnitude in the right cortex compared to the left cortex (Figure 4B). No early gamma power changes were observed in supplementary motor area or precentral gyrus. As reported from the z-score gamma power maps (Figure 2B), difference time courses showed limited signal changes at the early time points (Figure 4C) in agreement with the lack of statistically significant changes at early times in Figure 2B.

### 3.2. Late gamma power changes

For the combined trials, cluster-based permutation statistical analysis revealed that the significant early gamma power changes observed at <150 ms from stimulus onset increased in magnitude and spatial extent up to approximately 330 ms and then gradually diminished thereafter (Figure 2A and Figure 3A). However, considering the difference maps, the small statistically significant changes in gamma power observed at times > 150 ms continued to increase in magnitude and spatial extent at later timepoints (Figure 2B and Figure 3B). In particular, the difference maps showed that X-Press target trials showed greater gamma power in broad regions of the left hemisphere, which at later times coalesced into more focal sustained increases in the left precentral gyrus (button press was with the right thumb, Figure 3B). In addition, X-Press trials showed larger gamma power in the bilateral medial temporal parahippocampal gyri at ∼330 – 420 ms, and lower gamma power in bilateral anterior temporal cortex as well as in broad regions of the right hemisphere at later times (Figure 3B).

The ROI time courses displayed in Figure 4 confirmed the late gamma power findings observed in the cortical surface maps for times >150ms post-stimulus. In particular, for combined trials, the initial gamma power increases observed in bilateral caudal middle frontal, superior frontal, parahippocampal, parietal, occipital, and fusiform cortex during the 250 ms visual stimuli faded towards zero in the period after stimulus offset (Figure 4B). The difference analyses, on the other hand, showed delayed changes after stimulus offset in all regions, including greater left hemisphere gamma power for X-Press target trials in the left caudal middle frontal, superior frontal, parietal, occipital, fusiform and precentral cortex (traces above zero after stimulus offset in Figure 4B); as well as initially greater and then lower power changes for X-Press target trials in bilateral parahippocampal cortex (biphasic traces after stimulus offset in Figure 4B); and lower power for X-Press target trials at later times in right hemisphere in caudal middle frontal, superior frontal and parietal cortex (traces below zero after stimulus offset in Figure 4B).

## 4. Discussion

Using icEEG recordings, the current investigation aimed to study the gamma power changes associated with visual stimulus presentation and detection. Analyses focused on changes in gamma power at early and late periods post-stimulus presentation. The CPT paradigm used fully opaque letters that were displayed for 250 ms. Because these stimuli should be fully visible for participants, the earliest changes were hypothesized to represent electrophysiogical dynamics linked to conscious visual stimulus detection. The current results revealed a broad set of regions involved tens of milliseconds after stimulus presentation, comprising bilateral visual, prefrontal, medial temporal cortex, and left parietal and lateral temporal cortex. Meanwhile, at later times, sustained changes were observed overlapping the early response regions, but also recruiting more frontal, parietal, and temporal areas especially in the left hemisphere. These late responses may be linked to perceptual decision making – a process in which sensory information is integrated and used to produce motor responses. Likewise, for X-Press events, when a button press was made after the presentation of the target stimulus, relatively larger gamma power increases were observed in left motor and premotor regions associated with motor planning and execution; while relatively lower gamma power was observed in the right hemisphere.

### 4.1. Early gamma power changes

At early times, no statistically significant gamma power differences were observed between target (X-Press) and non-target (NonX-NoPress) events within 150 ms of stimulus onset (Figure 2B, 4C). This suggested that the neural responses in the gamma range at the very early stages of visual perception are similar regardless of the event type, allowing these events to be combined for further investigation. At the earliest time points post-stimulus in combined analyses, cortical surface maps (Figure 2A) and ROI time courses (Figure 4B) showed bilateral gamma power increases in prefrontal cortex, particularly in the caudal middle frontal cortex overlapping with FEF, orbital frontal cortex, superior and inferior frontal cortex. The rapidity of these responses suggests the dynamics in these first tens of milliseconds are involved in stimulus detection and sensory processing. Such findings are in line with previous human and non-human primate studies that suggest a major role for prefrontal cortex in the early stages of visual conscious perception within 100 ms from stimulus onset. In both human and non-human primate studies, early activity was observed within the prefrontal cortex (Panagiotaropoulos et al., 2012), especially in the FEF (Schmolesky et al., 1998; Blanke et al., 1999; Thompson and Schall, 1999, 2000; Bichot and Schall, 2002; Muggleton et al., 2003; O’Shea et al., 2004; Gregoriou et al., 2009; Libedinsky and Livingstone, 2011; Bollimunta et al., 2018). In another study, it was shown that sensory information can reach FEF within 50 ms, which is consistent with our findings (Kirchner et al., 2009; Kwon et al., 2021).

In addition to the gamma power changes associated with FEF, early gamma power bilateral increases were observed in orbital frontal cortex, superior and inferior frontal cortex (Figure 2A). A recent fMRI study reported that inferior frontal gyrus is involved in stimulus detection (Weilnhammer et al., 2021). The superior and orbitofrontal areas are thought to contribute to higher cognitive functions, but their particular role in detection requires further investigation (du Boisgueheneuc et al., 2006; Chaumon et al., 2014).

Early gamma power increases were observed in bilateral occipital and bilateral fusiform gyri (Figure 2A, 4B). These findings are consistent with previous human and non-human primate studies in visual stimulus detection (Schmolesky et al., 1998; Herman et al., 2019; Li et al., 2019). However, the early decrease observed in right anterior medial occipital cortex is more challenging to interpret. Two previous icEEG studies from our research group found suppression of gamma power in visual cortex during active task phases of the CPT task and after stimulus perception in a visual, perceptual threshold task (Herman et al., 2019; Li et al., 2019). These responses were speculated to act in increasing signal to noise during the CPT task active phase to improve performance and temporally suppress or focus subsequent stimulus processing. Similar explanations may be applicable for the current findings.

In another study, human medial temporal lobe and medial frontal cortex formed a target detection network during visual search, which corresponds with our findings (Figure 2A) (Wang et al., 2018). Moreover, the involvement of the medial temporal cortex may reflect its role higher-order stimulus processing and encoding (Hassabis et al., 2007; Schacter et al., 2007; Konkel and Cohen, 2009). The left lateral temporal-parietal cortex showed early gamma power increases (Figure 2A), suggesting that it might be a part of an early visual detection network. Prior work has shown that the left inferior occipitotemporal cortex is involved in processing of written words and letter strings, within 200 ms after stimulus presentation (Tarkiainen et al., 1999; Vinckier et al., 2007).

As hypothesized and anticipated by previous studies, we observed gamma power increases in the parietal cortex (Figure 2A, 4B). Previous studies suggested that the parietal cortex plays an important role in visual perception and attention (Critchley, 1962; Bisley and Goldberg, 2010). Specifically, the parietal cortex provides top-down attentional feedback to sensory networks that can lead to modulation of neuronal activity relevant to the early stages of sensory signal processing (Corbetta and Shulman, 2002; Saalmann et al., 2007; Gregoriou et al., 2009; Bisley and Goldberg, 2010). The critical role of the parietal cortex attention network is supported by clinical data which shows that damage of the parietal cortex can lead to spatial neglect (Parton et al., 2004).

### 4.2. Late gamma power changes

The gamma power increases observed in prefrontal, parietal, medial, and lateral temporal regions at the early stages after stimulus presentation persisted into late time periods (>150 ms) (Figure 2A and Figure 3A), suggesting that these regions may play important roles in both detection and higher-order processing, such as perceptual decision making (Hauser and Salinas, 2014). Numerous studies have identified frontoparietal networks involved in perceptual decision-making process (Kable and Glimcher, 2009; Li et al., 2009; Siegel et al., 2011; Mulder et al., 2012; Keuken et al., 2014; Yeon et al., 2020). In line with our findings (Figure 2A and Figure 3A), frontal areas of perceptual decision-making networks reported in the literature included inferior frontal gyrus, middle frontal gyrus, superior frontal gyrus, and orbitofrontal cortex. Our observed gamma power increases in the parietal cortex align with studies that reported intraparietal sulcus and inferior parietal lobule as a part of a frontoparietal network. Indeed, activity within the lateral intraparietal cortex has been shown to reflect accumulation of sensory evidence prior to response (Shadlen and Newsome, 2001). The changes observed in the anterior cingulate cortex (ACC) (Figure 2A) correspond with prior studies suggesting that the ACC may be involved in guiding decision making and providing the appropriate target-specific response (Thielscher and Pessoa, 2007; Scheibe et al., 2010; Yeon et al., 2020).

Widespread gamma power differences were observed in the left hemisphere for X-Press versus NonX-NoPress events between 330 ms and 510 ms in frontal, parietal and temporal areas (Figure 3B). These differences receded later leaving a persistent gamma power difference in SMA and precentral gyrus responsible for motor planning and execution. The gamma power increases associated with X-Press events in precentral gyrus are in line with prior work by Muthukumaraswamy (2010). Simultaneously, a persistent gamma power difference in the left medial and lateral temporal cortex was observed with relative gamma power decreases for X-Press target trials. Frontal, parietal, and temporal regions within the right cortex showed a sustained gamma power negative difference with a relative decrease for X-Press target trials (Figure 3B). The right cortex negative differences are in line with prior studies suggesting a right hemisphere dominance for response inhibition (Swick et al., 2011; Coxon et al., 2016).

Future investigations should compare the detection of responses across sensory modalities to determine if each sensory system involves unique spatiotemporal characteristics. Moreover, recruiting patients with subcortical depth electrodes could produce a powerful dataset to find the subcortical contributions along with the human detection cortical network described in this study. These additional studies will help to advance the investigation of the neural mechanism of detection, further elucidating the earliest stages of sensory processing. These findings may also provide additional insights into how sensory signals are later promoted for higher-order processing and implemented in complex behaviors.

## Acknowledgments

This work was supported by the Betsy and Jonathan Blattmachr Family and by the Loughridge Williams Foundation.

